# Function and structure of FlaK, a master regulator of the polar flagellar genes in marine *Vibrio*

**DOI:** 10.1101/2022.08.19.504623

**Authors:** Michio Homma, Tomoya Kobayakawa, Yuxi Hao, Tatsuro Nishikino, Seiji Kojima

## Abstract

*Vibrio alginolyticus* has a flagellum at the cell pole, and the *fla* genes, involved in its formation, are hierarchically regulated in several classes. FlaK (also called FlrA) is an ortholog of *Pseudomonas aeruginosa* FleQ, an AAA+ ATPase that functions as a master regulator for all later *fla* genes. In this study, we conducted mutational analysis of FlaK to examine its ATPase activity, ability to form a multimeric structure, and function in flagellation. We cloned *flaK* and confirmed that its deletion caused a non-flagellated phenotype. We substituted amino acids at the ATP binding/hydrolysis site and performed putative subunit interfaces in a multimeric structure. Mutations in the aforementioned sites abolished both ATPase activity and the ability of FlaK to induce downstream flagellar gene expression. The L371E mutation, at the putative subunit interface, abolished flagellar gene expression but retained ATPase activity, suggesting that ATP hydrolysis is not sufficient for flagellar gene expression. We also found that FlhG, a negative flagellar biogenesis regulator, suppressed the ATPase activity of FlaK. The 20 FlhG C-terminal residues are critical for reducing FlaK ATPase activity. Chemical crosslinking and size exclusion chromatography revealed that FlaK mostly exists as a dimer in solution and can form multimers, independent of ATP. However, ATP induced the interaction between FlhG and FlaK to form a large complex. The *in vivo* effects of FlhG on FlaK, such as multimer formation and/or DNA binding, are important for gene regulation.

**IMPORTANCE:** FlaK is an NtrC-type activator of the AAA+ ATPase subfamily of σ^54^-dependent promoters of flagellar genes. FlhG, a MinD-like ATPase, negatively regulates the polar flagellar number by collaborating with FlhF, an FtsY-like GTPase. We found that FlaK and FlhG interact in the presence of ATP, to form a large complex. Mutational analysis revealed the importance of FlaK ATPase activity in flagellar gene expression and provided a model of the *Vibrio* molecular mechanism that regulates the flagellar number.

## INTRODUCTION

Motile bacteria generally have flagella on their cell surface, which are rotated by a motor at their bases, enabling them to swim or swarm. The flagella number and position vary depending on the bacteria type and what it requires to achieve efficient movement within its habitat (1). Some bacterial species, such as *Pseudomonas* and *Vibrio* (*V. alginolyticus* and *V. cholerae*), have one flagellum at the cell’s pole. Genes involved in flagellar biogenesis are clustered in the chromosome, and their expression is collectively controlled as a flagellar regulon. These genes are assigned to three or four transcriptional hierarchies and sequentially expressed to construct the polar flagella. Marine bacterium of *V. alginolyticus*, used in this study, has four transcriptional hierarchies (Fig. S1A). The first class contains a single gene encoding a master regulator, FlaK, that has the FleQ and FlrA orthologs in *P. aeruginosa* and *V. cholerae*, respectively. These three master regulators are highly homologous (Fig. S2B).

The two genes, *flhF* and *flhG*, control the number and location of flagella that form at the cell pole (2–4). It has been shown that excessive FlhF levels in *V. alginolyticus* results in multiple flagella forming at the cell pole; conversely, its absence results in no flagellation, indicating that FlhF positively controls the number of flagella. Contrarily, when FlhG levels are increased, the cell does not form flagella, while its deficiency results in multiple flagella forming at the cell pole, demonstrating that FlhG negatively controls the flagella number (3). We demonstrated that ATPase activity and polar localization of FlhG was crucial for the ability to downregulate the number of polar flagella (5). The FlhF GTPase activity is stimulated by FlhG interaction (6). Furthermore, *P. aeruginosa* FleN, the ortholog of FlhG, acts on FleQ (the ortholog of FlrA or FlaK), and suppresses the expression of polar flagellar genes (7). Based on these insights, we had proposed a model for flagellar number control in *V. alginolyticus* (Fig. S1B) (6).

Among the flagellar master regulators, *P. aeruginosa* FleQ, is the best studied among the species (8, 9). It is a bacterial enhancer-binding protein (bEBPs) that belongs to the AAA+ ATPase family. This regulator contains a receiver domain (REC) at the N-terminal region, a medial AAA+ ATPase/σ^54^ interaction domain, a helix-turn-helix DNA-binding domain at the C-terminus, and a c-di-GMP-binding domain that regulates ATPase activity (Fig. S2A) (7, 10–12). The AAA+ ATPase domain is composed of Walker A and B motifs, which are responsible for nucleotide binding and catalytic reactions, respectively. Mutations in the Walker A (K180A) and Walker B (D245A or E246Q) motifs of FleQ reduce ATPase activity (9). In addition, the crystal structure of the AAA+ domain has been elucidated; it is divided into an α/β (SD1) and an α-helical subdomain (SD2) to form a ring-shaped hexamer structure (Fig. S2C) (9, 13). The FleQ mutant T149E and V380E, which corresponds to the boundary surface of the AAA+ -HTH structure (SD1-SD2) and the adjacent AAA+ domain (SD2-SD2), respectively, reduce the binding of c-di-GMP; however, the hexamer structure of FleQ is maintained and its ATPase activity, not reduced. Additionally, the introduction of mutant I374E, which inhibits the formation of hexamer rings, does not change the c-di-GMP binding capacity, but reduces ATPase activity.

The *V. alginolyticus* FlaK is composed of 488 amino acids, and has a molecular weight of ca. 55 kDa, with 54% sequence identity to FleQ (Fig. S2B). However, little is known about its function and structure. In this study, we examined whether FlaK functions in an ATPase activity-dependent manner, whether it adopts a multimeric structure, and whether FlhG, which negatively regulates flagellar number, suppresses FlaK function.

## RESULTS

### Properties of the constructed FlaK mutants

FlaK belongs to class 1 within this hierarchy (Fig. S1A). As described in the Introduction, FlaK is an AAA+ ATPase family protein, and its ATPase activity is likely related to its function as an σ^54^-dependent transcriptional activator. To test this hypothesis, we generated mutants based on the FlaK ortholog FleQ (7, 9). Based on sequence alignment (Fig. S2B), we constructed six mutants of FlaK; Q140E, K171A D236A, E237Q, I365E, and L371E. These correspond to sites of FleQ, T149E, K180A, D245A, E246Q, I374E, and V380E, respectively. The mutated residues were predicted to be located in the following functional sites: K171, the ATP binding site (Walker A motif); D236 and E237, the catalytic site for ATP hydrolysis (Walker B motif); I365, the subunit interface of the putative hexameric ring; Q140 and L371, the subunit interface of the c-di-GMP bound hexamer structure. The locations of these FlaK mutation sites are shown in the corresponding structures of FleQ (Fig. S2C).

The generated FlaK mutants were expressed in the arabinose-inducible plasmid pTSK146 in the *V. alginolyticus* Δ*flaK* strain (NMB362). We analyzed the bacterial motility. The expressed FlaK contained a hexa-histidine (His)-tag at the C-terminus. The Δ*flaK* strain NMB362 with the empty pBAD33 vector was nonmotile (Fig. 1A), whereas NMB362 expressing FlaK (with a His-tag) was motile, indicating that the His-tag did not affect FlaK function. The K171A, D236A, E237Q, I365E, and L371E mutations prevented motility; these include the three mutations predicted to disrupt ATP binding and hydrolysis. Only FlaK Q140E, predicted to be in the subunit interface of the c-di-GMP-bound hexamer, facilitated motility similar to that in the wild-type.

**Fig. 1.**
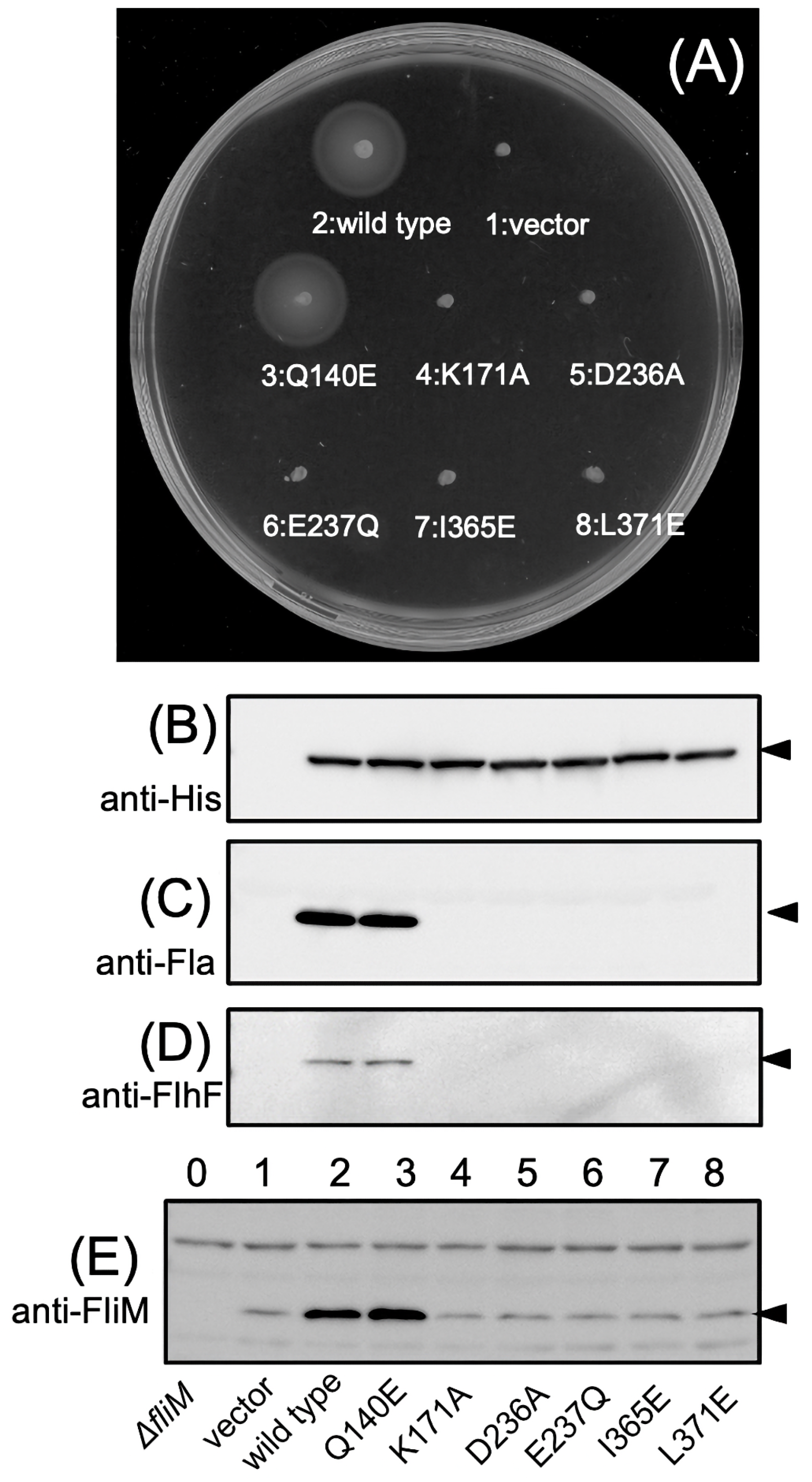
The *flaK* mutant cell motility and protein expression. (A) NMB362 (the *flaK* deletion strain) introduced with a pBAD33 vector plasmid (1: vector), pTSK146 encoding wild-type *flaK* (2: wild-type) or the pTSK146 derivative plasmids (Q140E, K171A, D236A, E237Q, I365E, L371E) containing the *flaK* mutant was cultured overnight at 30°C and aliquots of the overnight culture was spotted on the VPG agar plate containing arabinose and incubated at 30°C for 5 h. (B-E) The strains used in panel (A) were cultured at 30°C for 4 h in VPG medium containing arabinose. The proteins of whole cells were separated by SDS-PAGE and detected by western blotting using anti-Histidine (B), anti-Flagellin (C), anti-FlhF (D), or anti-FliM (E) antibody. Arrows indicate the detected proteins.

Next, we investigated the effects of FlaK mutations on the polar flagellar protein expression. Whole-cell samples of the Δ*flaK* strain expressing various mutant FlaK proteins were subjected to western blotting with antibodies against the His tag, FlhF, FliM, and flagellin (Fig. 1C-E). FlhF and FliM are expressed by class 2 genes, and flagellins are expressed by either class 3 or 4 genes. His-tagged wild-type and all FlaK mutants were detected at 55 kDa (Fig. 1B, lane 2-8). Both FlhF and FliM was detected in the wild-type and Q140E mutant motile strains (Fig. 1D, lanes 2 and 3) but was absent in the five nonmotile strains. Since no FliM (ca. 40 kDa) could be detected in the Δ*fliM* strain (NMB321) (Fig. 1E, lane 0), we conclude that a FlaK-independent promoter in the Δ*flaK* strain expresses low FliM levels.

Flagellins were only detected in strains expressing either wild-type or Q140E FlaK (Fig. 1C, lanes 2 and 3). This demonstrates that FlaK mutations disrupt ATP binding and hydrolysis (K171A, D236A, E237Q) and multimer formation (I365E, L371E), impairing the expression of class 2, 3, and 4 flagellar genes. Thus, these properties are important for FlaK function. Only the FlaK Q140E mutation had no effect on downstream flagellar protein expression, demonstrating that this mutation did not disrupt FlaK function, suggesting that this residue’s function may differ in the ortholog FleQ.

### FlhG modulates FlaK ATPase activity

To evaluate the multimeric state and ATPase activity of FlaK, we purified wild-type and six mutant FlaK proteins (Fig. S3), and examined their ability to hydrolyze ATP (Fig. 2A). The wild-type and Q140E mutant FlaK exhibited similar ATP hydrolysis activity. In contrast, the K171A, D236A, E237Q, and I365E FlaK mutants showed reduced ATPase activity. The L371E mutant hydrolyzed ATP at similar levels to that of the wild-type FlaK strain, demonstrating this mutation does not affect ATP hydrolysis.

**Fig. 2.**
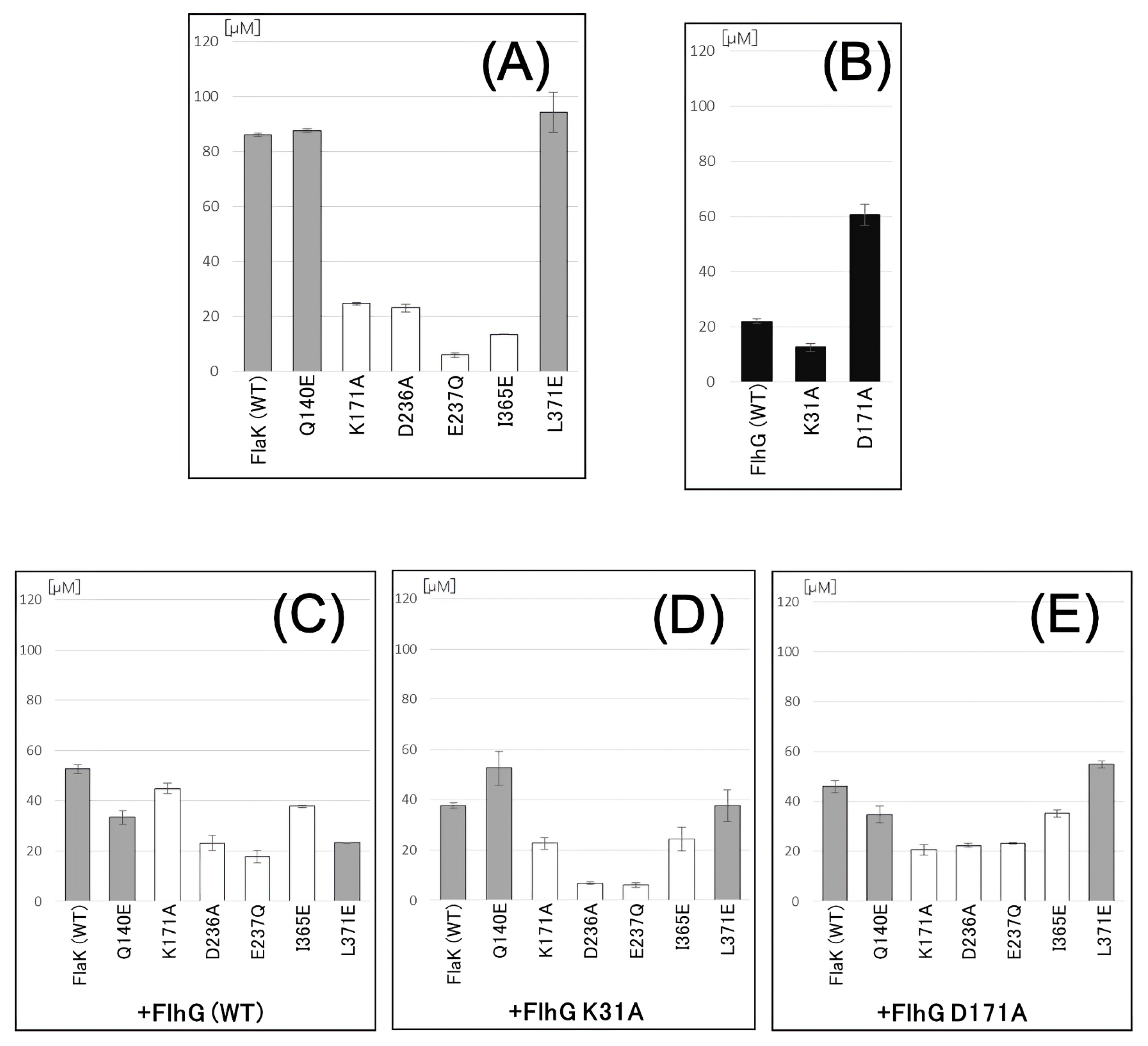
FlaK ATPase activity. The wild-type and mutant FlhG or FlaK proteins were prepared by His-tag affinity purification from *E. coli* cells harboring a plasmid pTrc-*flhG* or pTSK140. The release of inorganic phosphoric acid was measured when FlaK (A), FlhG (B), FlaK and FlhG-WT (C), FlaK and FlhG-K31A (D), or FlaK and FlhG-D171A (E) was mixed with ATP and MgCl_2_, and reacted for 30 min at 25°C.

The ATPase activity of *Pseudomonas aeruginosa* FleQ is inhibited by FleN (ortholog of *V. alginolyticus* FlhG) (7). Therefore, we investigated the suspected inhibiting effect of FlhG on the *V. alginolyticus* FlaK ATPase activity. To this end, we utilized His-tagged wild-type FlhG as well as K31A and D171A FlhG mutants (Fig. S4) (5). FlhG, itself, has low ATPase activity; the K31A mutant lacks ATPase activity, whereas the D171A mutant exhibits high ATPase activity (Fig. 2B) (5). Therefore, the inorganic phosphate, released by ATP hydrolysis in the mixture, was derived from both FlaK and FlhG ATPase activities. As shown in Fig. 2, when FlaK and FlhG proteins were mixed at equal concentrations, the activity of the wild-type, Q140E, and L371E FlaK mutants were greatly reduced. Similarly, the FlhG K31A and D171A mutants decreased the overall ATPase activity (Fig. 2C-E), indicating that FlhG can inhibit the ATPase activity of FlaK.

Next, we investigated the FlhG region responsible for inhibition of FlaK ATPase activity (Fig. 3). FlhG is a MinD/ParA-type ATPase and has an N-terminal extension sequence (20 residues) that is not conserved in MinD. This extension contains an FlhF-binding site with a DQAxxLR motif. Q9A and ΔN20, the deletion of N-terminal extension, FlhG mutants are unable to enhance FlhF GTPase activity (14). However, the Q9A and ΔN20 FlhG mutants decreased FlaK ATPase activity, indicating that these residues are not important for this activity. In contrast, an FlhG mutant that lacks the C-terminal 20 residues (ΔC20), which is predicted to form an amphipathic α-helix and is known to associate with the membrane, was unable to reduce FlaK ATPase activity. Thus, the FlhG C-terminus modulates FlaK ATPase activity. Finally, ATPase activity was increased by addition of the FlhG D171A mutant.

**Fig. 3.**
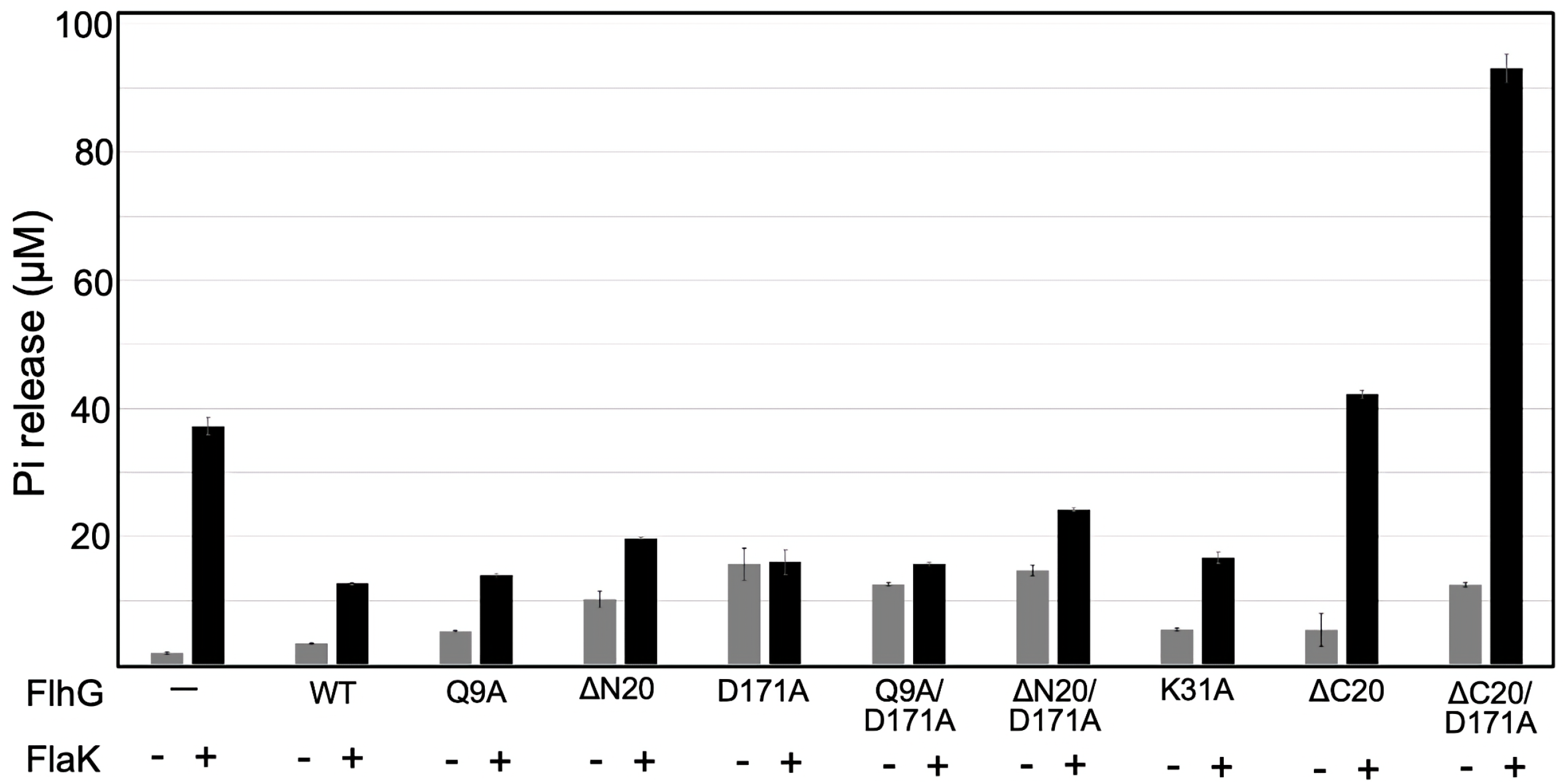
ATPase activity affected by FlhG mutants. The FlhG or FlaK proteins were prepared as Fig 2. The release of inorganic phosphoric acid was measured when FlaK and FlhG was mixed with ATP and MgCl_2_, and reacted for 30 min at 25°C.

### Conformational changes in FlaK

Structural changes in FlaK were first investigated using the EGS (Ethylene glycol bis (succinimidyl succinate)) chemical crosslinker which crosslinks amino groups. Both wild-type or mutant purified FlaK proteins were treated with EGS in the presence or absence of 1 mM ATP or non-hydrolyzable nucleotide AMP-PNP. The crosslinked products were separated by SDS-PAGE, followed by immunoblotting using anti-His antibody [detects FlaK; monomer putative calculated molecular weight (MW) of ca. 55 kDa] and is indicated by the white arrow (Fig. 4). The presumed dimer size (ca. 110 kDa), indicated by the black arrow, is apparent in all forms of FlaK, consistent with its dimer formation (see below). Larger crosslinked products > 180 kD (presumably tetramer, hexamer, and more), indicated by the gray arrow, were detected. Thus, indicating that FlaK can also form multimeric structures. The cross-linking profiles were not significantly affected by nucleotide addition. The E237Q and I365E mutants have relatively fewer multimeric forms than that of the other groups (Fig. 4).

**Fig. 4.**
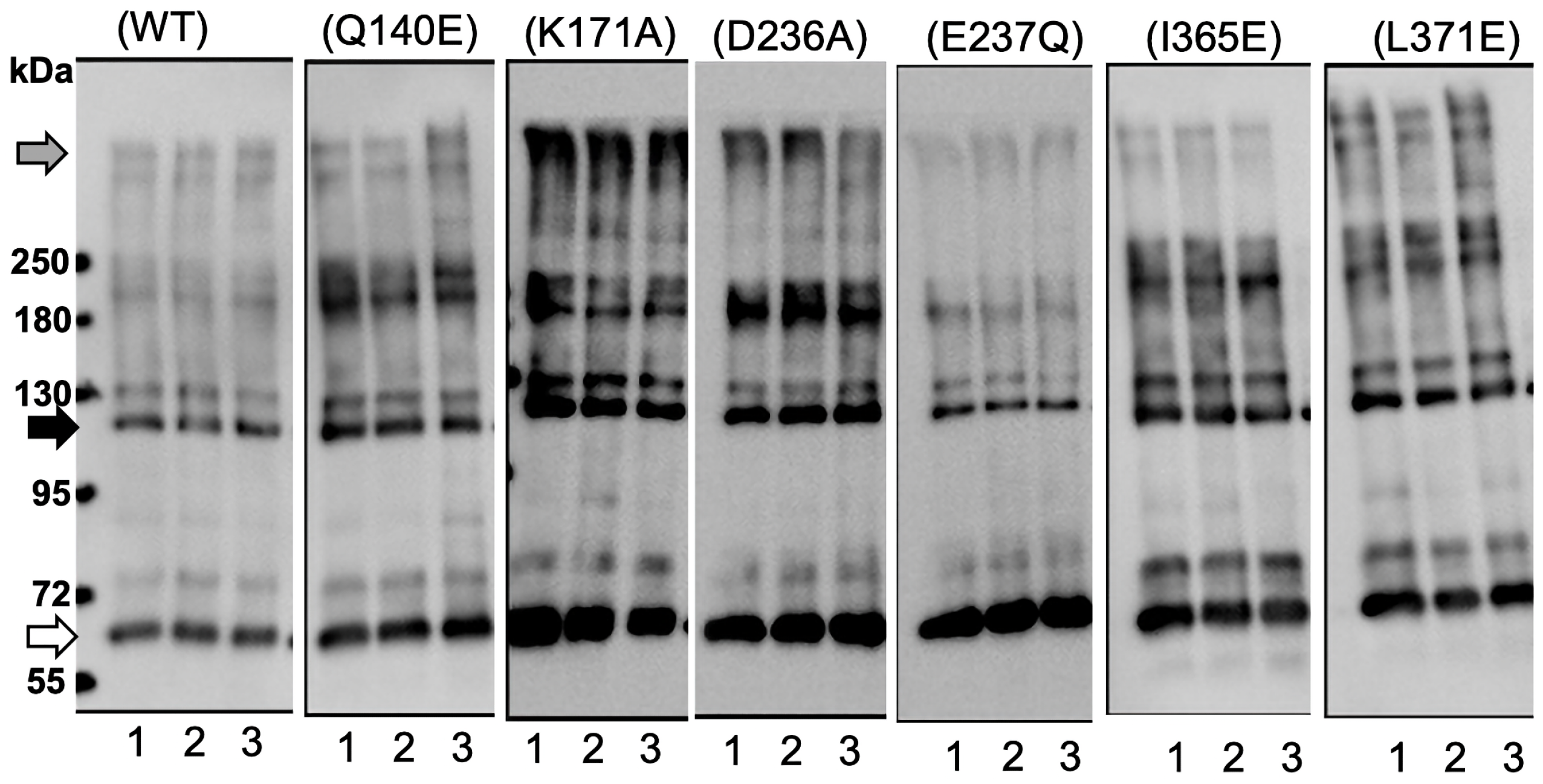
Cross-linking of FlaK. The purified proteins were treated with EGS in the presence of ATP (lane 3), AMP-PNP (lane 2) or absence of both (lane 1), and were separated by SDS-PAGE. The purified FlhG or FlaK proteins were prepared as Fig 2. The proteins were detected by western blotting using anti-His-tag antibody.

The oligomeric states of the wild-type, K171A, and I365E mutant FlaK proteins were analyzed by gel filtration. The wild-type and mutant FlaK proteins eluted at a MW of ca. 165 kDa, based on gel filtration size markers, regardless of the presence or absence of ATP. These results indicate that the FlaK protein is generally present as a dimer in solution.

FlaK was cross-linked with EGS, and underwent gel filtration, which revealed that the majority of cross-linked FlaK also eluted at ca. 165 kDa, but a new peak was apparent at approximately 300 kDa (Fig. 5). Immunoblotting the eluted fractions with anti-His antibody suggested that the cross-linked wild-type and K171A FlaK mutants also formed tetramers and hexamers (black arrow and white arrowhead, respectively) (Fig. 5A and C). The I365E FlaK mutant did not exhibit these distinct higher-order forms (Fig. 5B). Our results support the hypothesis that FlaK forms multimers.

**Fig. 5.**
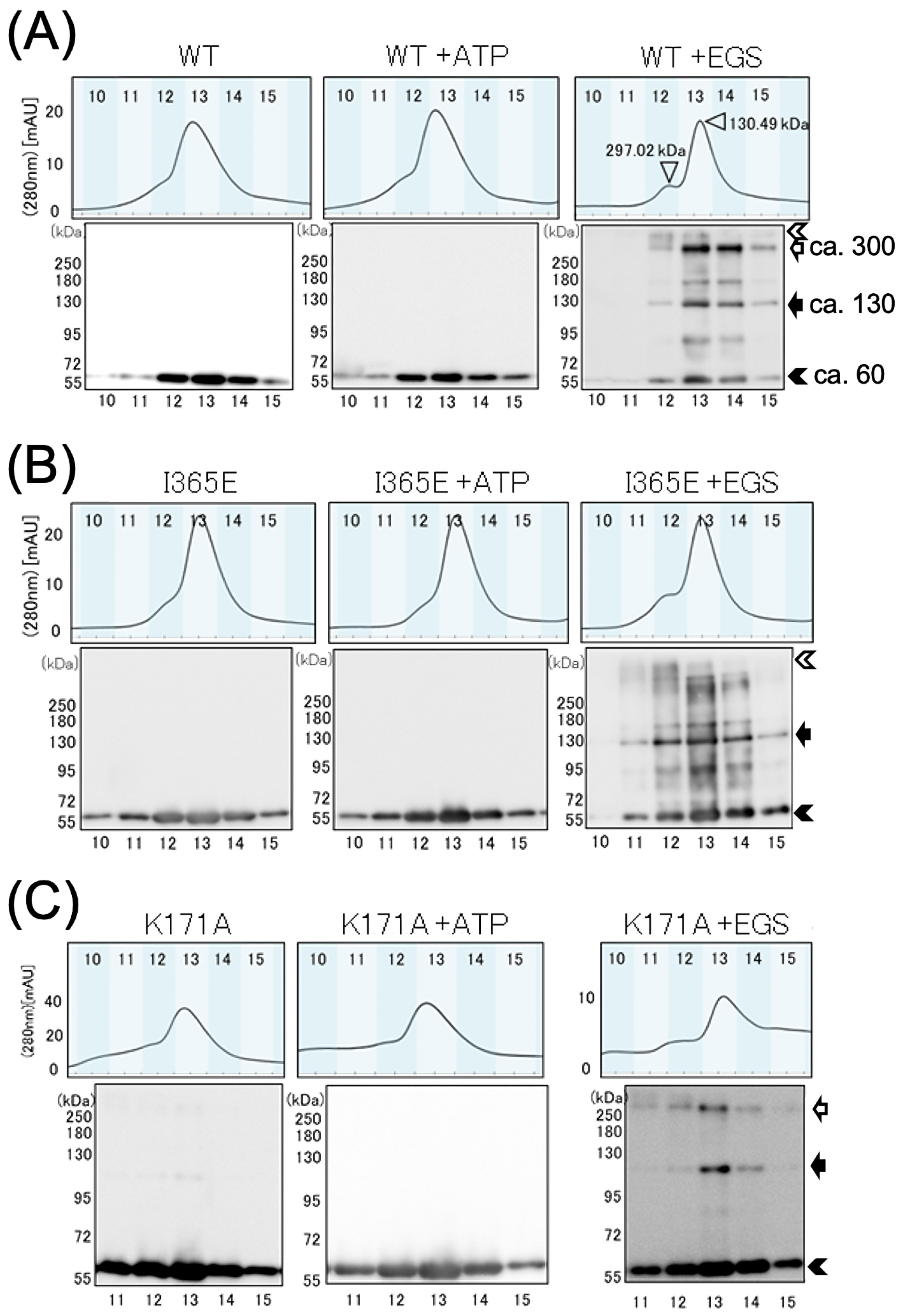
Analysis of cross-linking products of FlaK by size exclusion chromatography. After the purified proteins, wild-type (WT) FlaK (A), FlaK-I365E (B), and FlaK-K171A (C), were treated with ATP (+ATP) and ATP and EGS (+EGS), and the samples (500 μl), were injected into the size exclusion column (BIO-RAD ENrichTM SEC650) and eluted by buffer containing 1 mM ATP and 10 mM MgCl_2_. The elution profiles were monitored by the absorbance of 280 nm. The proteins in the fractions were separated by SDS-PAGE and detected by an immunoblot analysis, using anti-His-tag antibody (lower panels of elution profiles). The white and black arrows indicate the estimated sizes of the FlaK multimers. The black arrowhead indicates the FlaK monomer. We assume that the band indicated by white arrowheads corresponds to a hexamer eluted at the peaks of FlaK.

### FlaK and FlhG interaction

FlhG inhibited the ATPase activity of FlaK (see above), and gel filtration chromatography was used to determine whether FlhG and FlaK form a complex (Fig. 6). Both the wild-type and I365E mutant FlaK proteins were used, and they behaved similarly; the results are shown for the I365E FlaK protein. FlhG was mixed with I365E FlaK and subjected to gel filtration. The protein peaks corresponded to ca. 165 (FlaK) and ca. 29 kDa (FlhG). When 1 mM ATP was added to this sample and gel filtration was performed, the corresponding FlaK peak at 12.3 mL, was greatly reduced, and a peak appeared at 8.9 mL (Fig. 6C). This position corresponded to the void volume (>700 kDa). SDS-PAGE analysis of this peak region (Fig. 6D) showed that both FlaK and FlhG were present, supporting the idea that FlaK and FlhG interact to form large complexes.

**Fig. 6.**
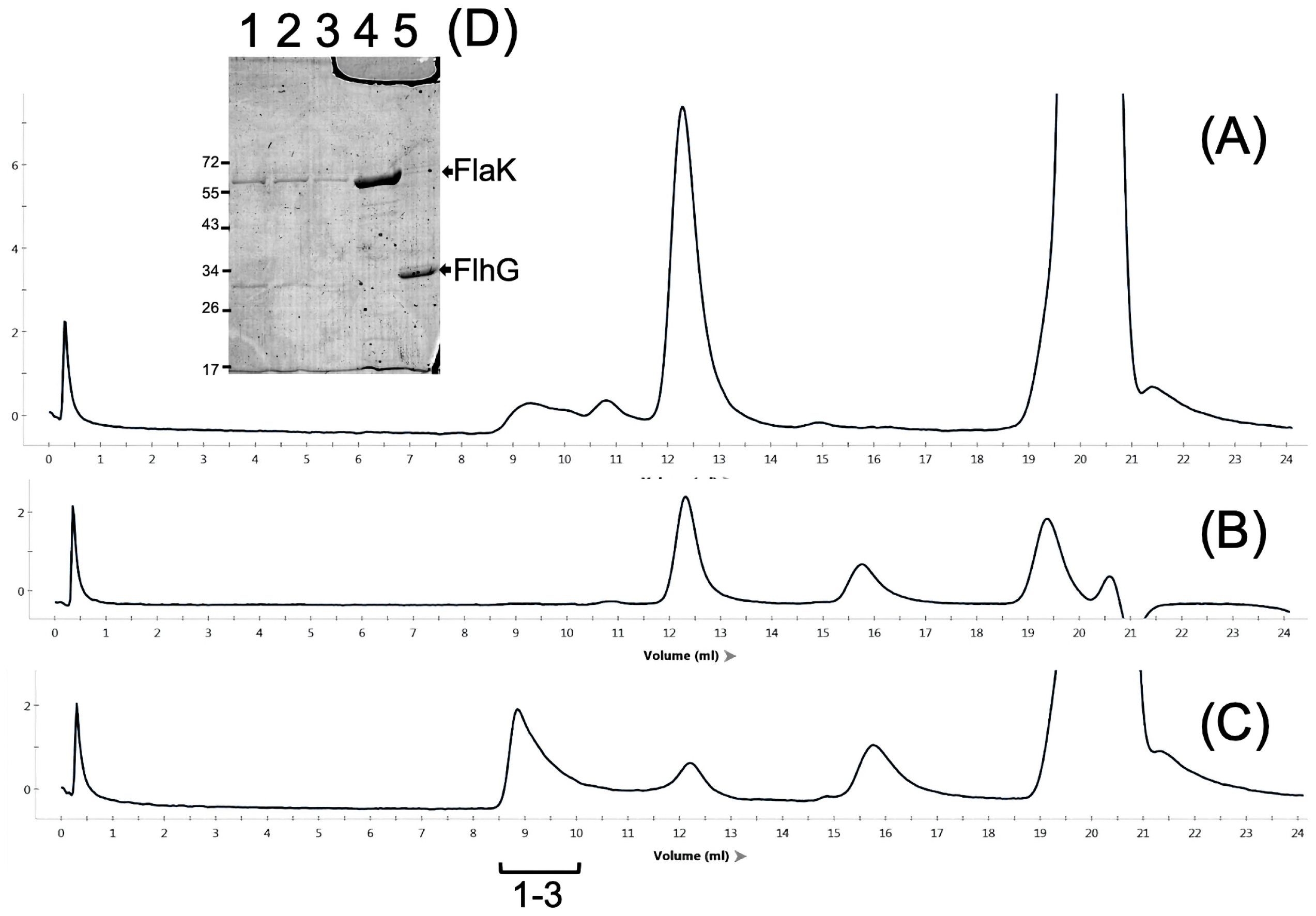
Interaction between FlaK and FlhG analyzed by size exclusion chromatography. The samples (100 μL) containing FlaK-I365E and 1 mM ATP (A), FlaK-I365E and FlhG (B), or FlaK-I365E, FlhG and 1 mM ATP (C) were injected into the size exclusion column (Superdex 200 Increase 10/300 GL) and eluted by 20TN150 buffer containing 5 mM MgCl_2_ and 10% glycerol. The elution profiles were monitored based on the absorbance at 280 nm. (D) Fractions were collected and proteins in the peak elution fractions, 18–20 (lanes 1-3), or the injected samples of FlaK-I365E (lane 4) and FlhG (lane 5) were analyzed with a 12% SDS-PAGE gel, followed by staining with Coomassie brilliant blue. The arrows indicate the band corresponding to FlaK (ca. 58 kDa) and FlhG (ca. 29 kDa).

Wild-type FlaK, with or without ATP, was observed via transmission electron microscopy. The size and shape of the FlaK protein varied, preventing an estimate of the number of molecules within the multimeric structure. ATP had no effect on these characteristics. Conversely, during gel filtration chromatography, when FlaK was mixed with FlhG in the presence of ATP, the proteins eluted at ca. 800 kDa and ca. 110 kDa. The large (>800 kDa) electron microscopy elution peak revealed the presence of large complexes (Fig. S5), indicating that FlaK and FlhG form a complex.

## DISCUSSION

The *Vibrio* regulation and biogenesis of polar flagella have been intensively studied (15). It is an excellent model system for investigating how the flagella location and number are determined. The genes involved in the formation and function of polar flagella are highly homologous across *Vibrio* spp., and have been studied in greater detail in *V. cholerae* (16), *V. parahaemolyticus* (17), and *V. campbellii* (18). Flagellin, the structural component of the flagellar filament, is the most abundant protein in the flagellum. In *Vibrio* spp., it has class 3 and 4 genes. Purified flagellin antibodies (likely composed of FlaABCD; referred to as PF45) have been generated in *V. alginolyticus* (19). Western immunoblotting with anti-PF45 antibodies confirmed that the five FlaK mutations, K171A, D236A, E237Q, I365E, and L371E, reduced the expression of class 3 and 4 genes, demonstrating the involvement of these residues in the flagellar hierarchy of *V. alginolyticus*. FliM protein levels, encoded by a class 2 gene, were also greatly reduced in the five mutants (Fig. S2, Fig. 1B, lane 2-8).

Based on homology with the *P. aeruginosa* flagellar master regulator FleQ (9, 13), three of the FlaK mutations, K171A, D236A, and E237Q (corresponding to K180A, D245A, and E246Q, respectively), are predicted to be involved in ATP binding and catalysis. The I365E FlaK mutation (corresponding to I374E of FleQ), is predicted to be involved in hexamer ring formation, a critical step in the stimulation of ATPase activity in σ^54^-dependent activators. FlaK is a σ^54^-dependent transcriptional activator and ATPase activity is required for transcriptional open complex formation by the σ^54^-RNAP holoenzyme. The low ATPase activity of these FlaK proteins, the lack of flagellin, and resulting non-motility of mutant strains are consistent with FlaK ATPase activity, which is critical for *V. alginolyticus* flagellar synthesis. Two FlaK mutations, Q140E and L371E (corresponding to T149E and V380E of FleQ, respectively), are localized to the hexamer interface border. These purified mutant proteins had ATPase activity similar to that of the wild-type. Interestingly, while strains expressing Q140E FlaK did not affect motility, those expressing L371E did not form polar flagella and were thus nonmotile (Fig. 1A, B). The V380E mutation in FleQ prevents multimer formation without affecting ATP hydrolysis. The hexameric FleQ ring structure is regulated by binding to c-di-GMP, which alters its conformation so it stimulates ATPase activity (9). We speculated that FlaK, in a manner similar to FleQ, forms a specific tight hexameric conformation that activates polar flagellar gene transcription.

FlhG (FleN), which controls flagellar number, interacts with FlaK (FlrA/FleQ) in *P. aeruginosa* and *S. putrifaciens* to downregulate flagellar synthesis. *S. putrifaciens* ATP-bound FlhG interacts with the HTH region of FlrA, which stimulates FlhG ATPase activity and downregulates FlrA-dependent class 2 gene transcription (20). When *V. alginolyticus* FlaK and FlhG were mixed, the ATPase activity of wild-type FlaK decreased because FlhG activity was very low. FlhG mutants with either no ATPase (K31A) or high ATPase (D171A) activity were able to suppress FlaK ATPase activity, suggesting that FlhG ATPase activity is not necessarily required for the suppression of FlaK transcriptional activity. Importantly, FlhG lacking C-terminal 20 residues, predicted to form an amphipathic α-helix, is unable to suppress FlaK ATPase activity. This result is consistent with *that of S. putrifaciens* FlhG, which requires an interaction between its C-terminal α7 and the FlrA HTH region to suppress FlrA activity and hyper-flagellation (20). This is also consistent with the crystal structure of the AAA+ domain of FleQ in complex with FleN (13), which shows that FleN binding allosterically to the central domain prevents ATP binding to FleQ and remodels the σ^54^ contact region to prevent transcription activation. It is currently unclear whether *V. alginolyticus* FlhG binds to the central (AAA+) or C-terminal (HTH) domain of *V. alginolyticus* FlaK.

The interaction between *V. alginolyticus* FlhG and FlaK was also detected using size exclusion chromatography. FlaK itself exists mainly as a dimer in solution (ca. 130 kDa), and ATP did not affect the oligomeric state of the wild-type, I365E, or K171A mutant FlaK proteins (Fig. 4). However, when EGS was added, a small 300 kDa peak of wild-type and mutant FlaK was detected, which may correspond to a putative hexameric complex. Western immunoblotting also detected a faint band at the presumed hexameric size (Fig. 3), which was reduced in the I365E mutant, presumed to have a defective hexamer ring formation.

Based on our results, we proposed a model for the role of FlaK in the formation of the polar flagellum of *V. alginolyticus* (Fig. 7). FlaK forms stable complexes with mostly dimeric structures, but the tight hexameric complex is the active form that hydrolyzes ATP and stimulates σ^54^-dependent transcription of class 2 flagellar genes. The interaction of the C-terminus of FlhG with FlaK disrupts this activity. Since the K31A FlhG mutant, which is unable to hydrolyze ATP, is still able to inhibit FlaK ATPase activity, it appears that the inhibitory effect does not require ATP hydrolysis by FlhG. As mentioned above, it is unclear whether *V. alginolyticus* FlhG interacts with the central and/or C-terminal FlaK domains. Further research will determine how FlhG influences FlaK-dependent transcriptional activation; either by inhibiting the FlaK-DNA interaction and/or hexamer formation and ATPase activity to provide a negative feedback mechanism controlling class 2 flagellar gene expression.

**Fig. 7.**
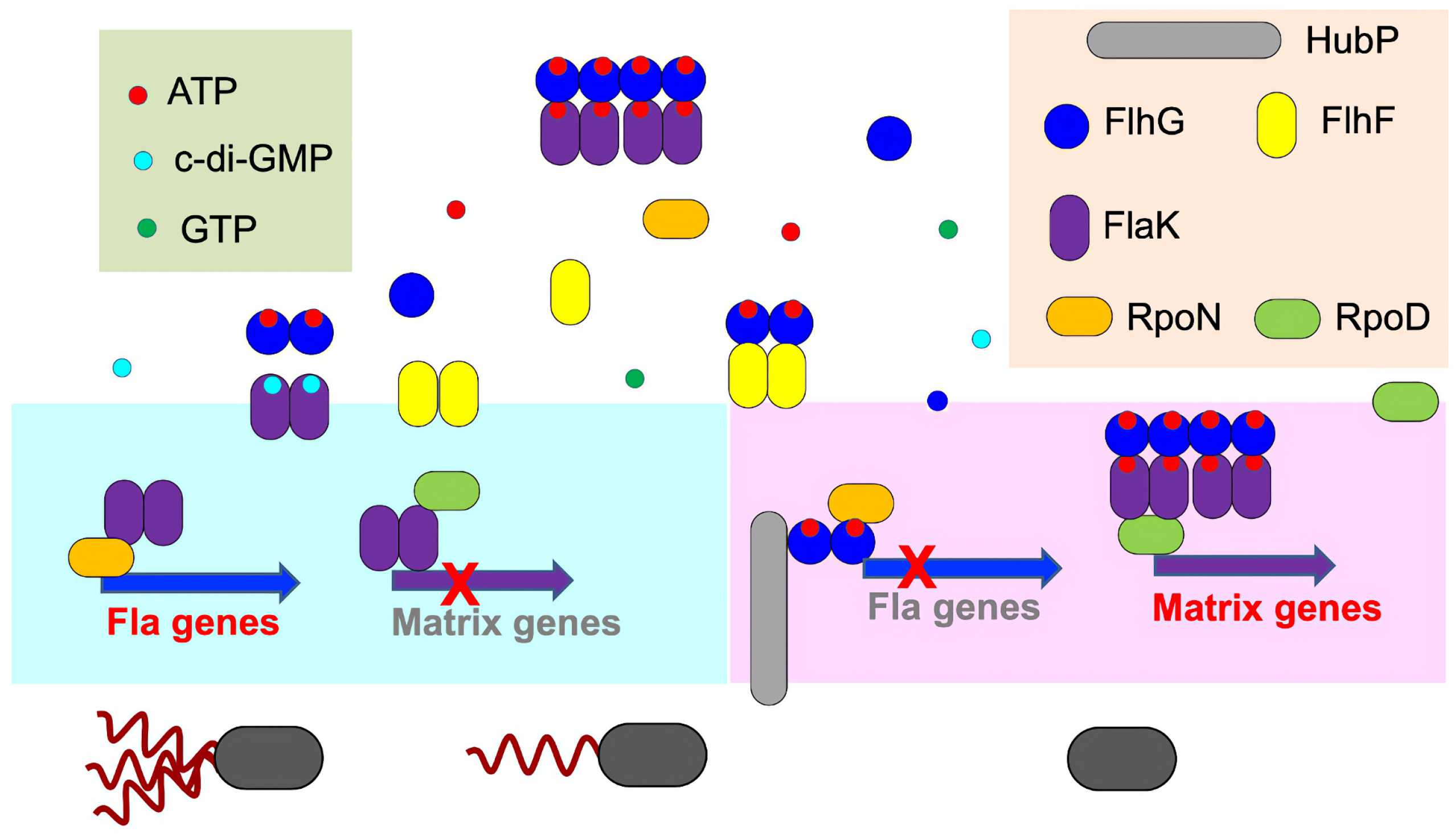
Working model of regulation for the polar flagella genes by FlaK. FlhG inhibits the function of FlaK and suppresses the formation of *Vibri*o polar flagella. The *rpoN* gene encodes the σ factor, σ54 (σ N, or RpoN), and the housekeeping σ factor, RpoD, and transcribes most genes in growing cells (8, 26, 27). HubP, a large membrane protein (ca. 160 kDa), is localized at cell poles to regulate the formation of flagella and pili, and interacts with various proteins, such as ParA and FlhG (28, 29).

## MATERIALS AND METHODS

### Strains used, culture medium and culture conditions

The strains and plasmids used are listed in Table S1. *E. coli* was grown in LB medium [1% (w/v) Bacto Tryptone, 0.5% (w/v) yeast extract, and 0.5% (w/v) NaCl] or SB medium [1.2% (w/v) Bacto Tryptone, 2.4% (w/v) yeast extract, 1.25% (w/v) K_2_HPO_4_, 0.38% (w/v) KH_2_PO_4_, and 0.5% (w/v) glycerol] at 37°C. To create a solid medium, agar was added at 1.25% (w/v) in LB medium. If necessary, chloramphenicol and ampicillin were added at final concentrations of 25 and 100 μg/ml, respectively.

*V. alginolyticus* was cultured at 30°C in the VC medium [0.5% (w/v) hipolypeptone, 0.5% (w/v) yeast extract, 0.4% (w/v) K_2_HPO_4_, 3% (w/v) NaCl, and 0.2% (w/v) glucose] or VPG medium [1% (w/v) hipolypeptone, 0.4% (w/v) K_2_HPO_4_, 3% (w/v) NaCl, and 0.5% (w/v) glycerol]. For growth on solid medium, agar was added to 1.25% (w/v) VC medium. A solid and soft agar media was prepared by adding 1.25% agar and 0.25% Bacto Agar, respectively. If necessary, chloramphenicol was added at a final concentration of 2.5 μg ml^−1^. The gene, under control of the arabinose promoter (pBAD), was induced by adding an amount of arabinose (0.02%) suitable for the expression level.

### DNA manipulations, mutagenesis and sequencing

Routine DNA manipulations were performed, according to standard procedures. Mutations in *flhG* were introduced into plasmids by the “QuikChange” site-directed mutagenesis method, as described by Agilent Technologies (Santa Clara, CA, USA); DNA sequencing confirmed each mutation. The *flaK* deletion strain, NMB362, was created by allelic exchange, using the suicide vector, pSW7848, and homologous recombination, as previously described (21, 22). Briefly, the *flaK* gene, with 250 bp up and downstream sequences, was PCR-amplified. Thereafter, a 967 bp internal *flaK* sequence was deleted and cloned into the pGEM-T easy vector to generate pTSK138. The internal deletion caused a FlaK frameshift at Pro84. The cloned sequence was transferred to the suicide vector, pSW7848, to generate pTSK138-2. We performed the conjugational transfer of pTSK138_2 from *E. coli* β3914 to the wild-type strain, for polar flagellar motility (VIO5). Thereafter, we carried out an allelic exchange, as described previously (22). As previously described, *Vibrio* was transformed by electroporation, using a Gene Pulser from Bio-Rad (23).

### Immunoblotting

The protein solution was mixed with SDS loading buffer and boiled at 95°C for 5 min. Proteins were separated by SDS-PAGE and then transferred onto polyvinylidene difluoride (PVDF) membranes. The primary antibodies anti-His-tag antibody (MBL Co. LTD), anti-FlhF antibody (24), anti-FliM antibody (FliM B0549, this study), and anti-PF45 antibody (19) detected FlaK, FlhF, FliM, and flagellin. HRP-labeled rabbit anti-IgG antibody (Thermo Scientific) was the secondary antibody. Using the ECL reaction solution, the proteins were detected with LAS 3000 (Fujifilm).

### Antibody against FliM

*E. coli* overproduced a recombinant FliM that was purified and used as an antibody against *V. alginolyticus* polar FliM. For the overproduction of FliM, the plasmid pMK2001 (pET15b based plasmid encoding an N-terminal His-tagged FliM) was introduced into *E. coli* BL21(DE3). Overproduction and purification of His-tagged FliM were performed as previously described (25). Purified FliM (3 mg) was separated by SDS-PAGE. The FliM band was excised and used to inoculate the rabbit protein samples. Rabbit anti-FliM antibody (FliM B0549) was produced by Biogate Co.

### FlaK protein purification

*E. coli* BL21(DE3) was transformed with pTSK140 (encoding wild-type Flak with a C-terminal His_6_-tag) and its derivatives (encoding mutant *flaK*). The cells were inoculated in 1 L of LB medium with ampicillin (final concentration, 100 μg ml^−1^) and incubated with shaking (150 rpm at 37°C) until the OD_660_ nm value reached 1.0. We then added IPTG (final concentration 0.4 mM), and the flask was chilled with ice water to induce protein expression. After the cells were re-incubated with shaking for 3 h at 18°C, they were centrifuged at 3,500 × *g* for 10 min at 4°C, and the cell pellet was stored at −80°C.

The stored cells were thawed and suspended in 40 mL of 20TN150 buffer (20 mM Tris-HCl, pH 8.0, 0.15 M NaCl) that contained 5 mM imidazole and 10 mg lysozyme. Cells were crushed by sonication (power = 8, duty = 50% for 1 min, 5 times) on ice, and the uncrushed bacteria were removed by low-speed centrifugation (3,500 × *g* for 10 min at 4°C). The supernatant was separated by Himac CP80WX ultracentrifugation (154,000 × *g* for 30 min at 4°C), using the P50AT2 rotor. The soluble fraction was loaded onto Ni-NTA resin (2 ml), equilibrated with 20TN150 containing 5 mM imidazole. After 10 repeated washings with the 20TN150 volume containing 60 mM imidazole, the proteins were eluted with 20TN150 containing 300 mM imidazole. The resulting FlaK solution was loaded onto a desalting column, MidiTrap G-25 (Cytiva), with 20TN150. The obtained FlaK was frozen in liquid nitrogen and stored at −80°C. A spectrophotometer measured the absorbance of the purified FlaK protein; the A280 and A320 values and the absorption coefficient was used to calculate the protein concentration.

### Purification of FlhG protein

*E. coli* BL21(DE3)/pLysS was transformed with pTrc-*his-tev-flhG* (encoding wild-type FlhG with a His-tag) or its derivatives (encoding mutant FlhG). Cells were grown in 1 L of SB medium containing chloramphenicol and ampicillin with a final concentration of 25 and 100 μg/mL, respectively, and incubated with shaking (150 rpm at 37°C) until the OD_660nm_ = 1.0. IPTG (final concentration 0.4 mM) was then added, and the flask was chilled with ice water. The cells were re-incubated for 3 h at 18°C, centrifuged at 3,500 × *g* for 10 min at 4°C, and stored at −80°C.

The cells were thawed, and 40 mL of 20TN300 buffer (20 mM Tris-HCl, pH 8.0, 0.3 M NaCl) containing 10% glycerol and PMSF (final concentration 0.5 mM) was added. The cells were disrupted by sonication (power = 8, duty = 50%, 1 min, 5 times) on ice, and the undisrupted bacteria were removed by low-speed centrifugation (3,500 × *g* for 10 min at 4°C). The soluble fraction was then separated by ultracentrifugation (154,000 × *g* for 30 min at 4°C) and passed through a HiTrap TALON column (5 mL), equilibrated with Buffer E. The column, with adsorbed FlhG, was connected to an AKTA prime system (Cytiva) and washed with 20TN300 containing 10% glycerol and 20 mM imidazole, and eluted with a buffer containing 65 mM imidazole. During elution, 1 mL fractions were collected, and the peak fractions were frozen in liquid nitrogen and stored at −80°C.

### Measurement of ATPase activity

The ATPase activity of the FlaK and FlhG proteins, mixed with ATP using a kit from Innova Bioscience, was evaluated by measuring the inorganic phosphate concentration. The 96-well plates were prepared with 50 μl of the protein solution (a final concentration of 2.5 μM of each protein) and 50 μl of 20TN150, containing 10 mM MgCl_2_ and 1 mM ATP. These reagents were allowed to react for 30 min at 25°C. Next, 25 μl of gold mix, a 100:1 mixture of PiColorLock Gold and Accelerator, was added to stop the ATP hydrolysis reaction. Then, 10 μl of stabilizer was added to the sample, which was re-incubated at 25°C for 30 min. The 595 nm absorbance of the sample was measured using a plate reader. The results were plotted on a calibration curve showing the relationship of the phosphate solution absorbance and gold mix at different concentrations (0, 10, 20, 50, and 100 μM) at 595 nm. The concentration of inorganic phosphate was calculated using the curve. Each sample was measured three times, and the average value was calculated.

### Cross-linking reaction

Ethylene glycol bis (succinimidyl succinate) (EGS) was used for cross-linking FlaK. Immediately after purification, the FlaK protein solution was replaced with HEPES buffer (20 mM HEPES, 0.15 M NaCl) using a desalting column MidiTrap G-25, and the concentration of FlaK was adjusted to 5 μM. The reagents MgCl_2_, ATP or AMP-PNP, and EGS were added to the solution to achieve respective concentrations of 10 mM, 1 mM, and 0.25 mM. The reaction was carried out at 25°C for 30 min, after which 20 mM Tris-HCl (pH 8.0) was added to stop the EGS reaction.

### Gel filtration chromatography of FlaK proteins after the treatment of EGS

Immediately after purification, 15 μM of the FlaK protein solution was prepared in HEPES buffer. When the concentration of the purified protein was <15 μM, it was used without dilution. The reaction with ATP was adjusted to a final concentration of 1 mM ATP and reacted at 25°C for 30 min. The reaction with EGS was adjusted to the final concentration of 0.25 mM and reacted for 30 min at 25°C. After the aggregates were removed by ultracentrifugation (154,000 × *g* for 30 min at 4°C), 500 μl of the supernatant was injected into a column (BIO-RAD ENrich™ SEC650) and run with either 20TN150 or HEPES buffer containing 1 mM ATP and 10 mM MgCl_2_, at a flow rate of 0.75 ml/min, and 1 ml of each fraction was collected. SDS samples were prepared from the collected fractions and detected by immunoblotting with anti-His antibody.

### Negative staining preparation for electron microscopy images

A 5 μl solution was applied to a glow-discharged continuous carbon grid. The excess solution was removed using filter paper, and the sample was subsequently stained with 2% ammonium molybdate on a carbon grid. The H-7650 transmission electron microscope (Hitachi), operated at 80 kV and equipped with a FastScan-F114 CCD camera (TVIPS, Gauting, Germany) recorded the results at a nominal × 80,000 magnification.

## ACKNOWLEDGMENTS

We thank Dr. Karl Klose for critically reading the manuscript, Dr. Kimika Maki for electron microscopy technical support, and Akiko Abe for constructing the plasmids and strains. This work was supported, in part, by JSPS KAKENHI: Grant Numbers; 16H04774 (to S.K.) and 20H03220 (to M.H.).

## SUPPLEMENTAL MATERIAL

The supplementary information associated with this article can be found in the online version of the publisher’s website.

## REFERENCES

1. Kojima S, Terashima H, Homma M. 2020. Regulation of the Single Polar Flagellar Biogenesis. Biomolecules 10:533.

2. Correa NE, Peng F, Klose KE. 2005. Roles of the regulatory proteins FlhF and FlhG in the *Vibrio cholerae* flagellar transcription hierarchy. J Bacteriol 187:6324–6332.

3. Kusumoto A, Kamisaka K, Yakushi T, Terashima H, Shinohara A, Homma M. 2006. Regulation of polar flagellar number by the *flhF* and *flhG* genes in *Vibrio alginolyticus*. J Biochem (Tokyo) 139:113–121.

4. Balaban M, Hendrixson DR. 2011. Polar flagellar biosynthesis and a regulator of flagellar number influence spatial parameters of cell division in *Campylobacter jejuni*. PLoS Pathog 7:e1002420.

5. Ono H, Takashima A, Hirata H, Homma M, Kojima S. 2015. The MinD homolog FlhG regulates the synthesis of the single polar flagellum of *Vibrio alginolyticus*. Mol Microbiol 98:130–141.

6. Kondo S, Imura Y, Mizuno A, Homma M, Kojima S. 2018. Biochemical analysis of GTPase FlhF which controls the number and position of flagellar formation in marine *Vibrio*. Sci Rep 8:12115.

7. Baraquet C, Harwood CS. 2013. Cyclic diguanosine monophosphate represses bacterial flagella synthesis by interacting with the Walker A motif of the enhancer-binding protein FleQ. Proc Natl Acad Sci U S A 110:18478–18483.

8. Bush M, Dixon R. 2012. The role of bacterial enhancer binding proteins as specialized activators of σ54-dependent transcription. Microbiol Mol Biol Rev 76:497–529.

9. Matsuyama BY, Krasteva PV, Baraquet C, Harwood CS, Sondermann H, Navarro MV. 2016. Mechanistic insights into c-di-GMP-dependent control of the biofilm regulator FleQ from *Pseudomonas aeruginosa*. Proc Natl Acad Sci U S A 113:E209–18.

10. Arora SK, Ritchings BW, Almira EC, Lory S, Ramphal R. 1997. A transcriptional activator, FleQ, regulates mucin adhesion and flagellar gene expression in *Pseudomonas aeruginosa* in a cascade manner. J Bacteriol 179:5574–5581.

11. Jyot J, Dasgupta N, Ramphal R. 2002. FleQ, the major flagellar gene regulator in *Pseudomonas aeruginosa*, binds to enhancer sites located either upstream or atypically downstream of the RpoN binding site. J Bacteriol 184:5251–5260.

12. Dasgupta N, Wolfgang MC, Goodman AL, Arora SK, Jyot J, Lory S, Ramphal R. 2003. A four-tiered transcriptional regulatory circuit controls flagellar biogenesis in *Pseudomonas aeruginosa*. Mol Microbiol 50:809–824.

13. Chanchal, Banerjee P, Raghav S, Goswami HN, Jain D. 2021. The antiactivator FleN uses an allosteric mechanism to regulate σ^54^-dependent expression of flagellar genes in P*seudomonas aeruginosa*. Sci Adv 7:eabj1792.

14. Homma M, Mizuno A, Hao Y, Kojima S. 2022. Functional analysis of the N-terminal region of *Vibrio* FlhG, a MinD-type ATPase in flagellar number control. J Biochem (Tokyo) 172:99–107.

15. Homma M, Nishikino T, Kojima S. 2022. Achievements in bacterial flagellar research with focus on *Vibrio* species. Microbiol Immunol 66:75–95.

16. Echazarreta MA, Klose KE. 2019. *Vibrio* Flagellar Synthesis. Front Cell Infect Microbiol 9:131.

17. McCarter LL. 2006. Regulation of flagella. Curr Opin Microbiol 9:180–186.

18. Petersen BD, Liu MS, Podicheti R, Yang AY, Simpson CA, Hemmerich C, Rusch DB, van Kessel JC. 2021. The polar flagellar transcriptional regulatory network in *Vibrio campbellii* deviates from canonical *Vibrio* species. J Bacteriol 203:e0027621.

19. Nishioka N, Furuno M, Kawagishi I, Homma M. 1998. Flagellin-containing membrane vesicles excreted from *Vibrio alginolyticus* mutants lacking a polar-flagellar filament. J Biochem (Tokyo) 123:1169–1173.

20. Blagotinsek V, Schwan M, Steinchen W, Mrusek D, Hook JC, Rossmann F, Freibert SA, Kratzat H, Murat G, Kressler D, Beckmann R, Beeby M, Thormann KM, Bange G. 2020. An ATP-dependent partner switch links flagellar C-ring assembly with gene expression. Proc Natl Acad Sci U S A 117:20826–20835.

21. Le Roux F, Binesse J, Saulnier D, Mazel D. 2007. Construction of a *Vibrio splendidus* mutant lacking the metalloprotease gene *vsm* by use of a novel counterselectable suicide vector. Appl Environ Microbiol 73:777–784.

22. Kojima S, Yoneda T, Morimoto W, Homma M. 2019. Effect of PlzD, a YcgR homologue of c-di-GMP-binding protein, on polar flagellar motility in *Vibrio alginolyticus*. J Biochem (Tokyo) 166:77–88.

23. Kawagishi I, Okunishi I, Homma M, Imae Y. 1994. Removal of the periplasmic DNase before electroporation enhances efficiency of transformation in a marine bacterium *Vibrio alginolyticus*. Microbiology 140:2355–2361.

24. Kondo S, Homma M, Kojima S. 2017. Analysis of the GTPase motif of FlhF in the control of the number and location of polar flagella in *Vibrio alginolyticus*. Biophys Physicobiol 14:173–181.

25. Homma M, Takekawa N, Fujiwara K, Hao Y, Onoue Y, Kojima S. 2022. Formation of multiple flagella caused by a mutation of the flagellar rotor protein FliM in *Vibrio alginolyticus*. Genes Cells. in press, doi:10.1101/2022.04.21.489128:2022.04.21.489128.

26. Österberg S, del Peso-Santos T, Shingler V. 2011. Regulation of alternative sigma factor use. Annu Rev Microbiol 65:37–55.

27. Feklístov A, Sharon BD, Darst SA, Gross CA. 2014. Bacterial sigma factors: a historical, structural, and genomic perspective. Annu Rev Microbiol 68:357–76.

28. Takekawa N, Kwon S, Nishioka N, Kojima S, Homma M. 2016. HubP, a polar landmark protein, regulates flagellar number by assisting in the proper polar localization of FlhG in *Vibrio alginolyticus*. J Bacteriol 198:3091–3098.

29. Altinoglu I, Abriat G, Carreaux A, Torres-Sánchez L, Poidevin M, Krasteva PV, Yamaichi Y. 2022. Analysis of HubP-dependent cell pole protein targeting in *Vibrio cholerae* uncovers novel motility regulators. PLoS Genet 18:e1009991.

